# Disrespect and abuse of women during childbirth in public health facilities in Arba Minch town, South Ethiopia – a cross-sectional study

**DOI:** 10.1101/430199

**Authors:** Gebresilasea Gendisha Ukke, Mekdes Kondale Gurara, Wanzahun Godana Boynito

## Abstract

**Introduction:** Disrespect and abuse during childbirth is the main deterring factor to skilled birth utilization as compared to other more commonly known factors such as financial and physical inaccessibility.

**Objective:** To assess the occurrence of women’s disrespect and abuse during childbirth in public health facilities in Arba Minch town, south Ethiopia.

**Methods:** Institution based cross-sectional study design was employed at all public health institutions in Arba Minch town, south Ethiopia. Systematic sampling method was used to include 281 women who had given birth at public health institutions between January 01 and February 28, 2017. Data were collected by face to face interview by four midwife tutors and supervised by the principal investigator on daily bases. Semi-structured pretested questionnaire was used to collect data. Epi info version 7.1.2.0 and SPSS version 24 were used to enter and analyze the data respectively.

**Results:** A total of 281 women were participated in this study. The overall prevalence of non-respectful care was 98.9%. The women’s right to information and informed consent was the most frequently violated right with a prevalence of 92.5% (95% CI: 90.9, 94.1) followed by non-dignified care (36.7, 95% CI: 34.9, 38.5), physical abuse (29.5%, 95% CI: 24.2, 34.8), discrimination (18.1%, 95% CI: 13.6, 22.6), non-confidential care (17.1%,95% CI: 12.7, 21.5) and abandonment of care (4.3%, 95% CI: 3.1, 5.5). However, there is no woman who had been kept in detention in the health facilities. Being rural resident, giving birth in hospital, having no or low educational status and giving birth by cesarean route were factors which were significantly associated with specific women’s rights violations.

**Conclusions and recommendations:** The status of non-respectful and abusive care in the health care facilities in this study area is unacceptably high and needs serious attention by the health managers to tackle the problem.

## Introduction

Childbirth is a special moment of happiness for women and their families but it is also a time of intense vulnerability. Women might not get their expected quality of care and level of respect by the health care providers. When they are treated disrespectfully, the negative encounter of women with health workers during delivery leaves long lasting damage and emotional trauma which can significantly affect skilled birth attendance negatively [1].

For the past several years, Safe Motherhood Initiative has focused mainly on improving access to and utilization of skilled childbirth attendance and facility-based maternity care. Increasing the proportion of women in developing nations who deliver with skilled attendance is being advocated as the single most important intervention to decrease maternal mortality and morbidity. However, client satisfaction with care which is an important element of quality care and hence influences effectiveness of care has not been given much attention. As research indicates, problems related to care provider behavior and attitude are more important deterring factors than geographical and financial limitations to utilization of skilled childbirth care[2, 3].

Respectful maternity care recognizes that safe motherhood must be expanded beyond the prevention of morbidity or mortality to encompass respect for women’s basic human rights, including respect for women’s autonomy, dignity, feelings, choices, and preferences, such as having a companion wherever possible. The respectful maternity care approach is centered on the individual and based on principles of ethics and respect for human rights. The Respectful Maternity Care Charter developed by the White Ribbon Alliance and respectful maternity care partners, is based on a framework of human rights and is a response to the growing body of evidence documenting disrespect for and abuse of childbearing women [1, 2, 4].

According to the charter, seven rights were drawn from the categories of disrespect and abuse which are not mutually exclusive: *Article 1:* Every woman has the right to be free from harm and ill treatment. *Article 2:* Every woman has the right to information, informed consent and refusal, and respect for her choices and preferences, including the right to her choice of companionship during maternity care. Article *3:* Every woman has the right to privacy and confidentiality. *Article 4:* Every woman has the right to be treated with dignity and respect.*Article 5:* Every woman has the right to equality, freedom from discrimination, and equitable care.*Article 6:* Every woman has the right to healthcare and to the highest attainable level.*Article 7:* Every woman has the right to liberty, autonomy, self-determination, and freedom from coercion[2].

Even though significant progress had been made in improving maternal health between 1990 and 2015, maternal morbidity and mortality still continues to be a public health problem globally. In 2015, an estimated 303,000 women worldwide have lost their life due to easily preventable pregnancy and childbirth related complications, 99% of which are contributed by low income countries [5].

Ethiopia is one of the countries with the highest maternal mortality ratio in the world. According to the latest (2016) Ethiopian demographic and health survey report, maternal mortality ratio of the country was 412 per 100, 000 live births. Though the country is moving ahead in reducing maternal mortality; to achieve the newly set sustainable developmental goal three: reducing the global maternal mortality ratio to less than 70,000 per 100,000 live births in 2030, tripling the former 2.3% annual maternal mortality reduction rate to 7.5% is expected[5–7].

Increasing skilled birth attendance has been one of the strategies proposed by experts to reduce the number of women who lose their life as a result of pregnancy and child birth, as majority of deaths occur during delivery and immediate postpartum periods [8, 9].

Although many years have been passed since the strategy was proposed, the proportion of women who are giving birth in health institutions (by trained birth attendants) in Ethiopia is not more than 15%. By this pace, the issue of achieving the sustainable development goal by 2030, will be questionable[5, 10]

Disrespect and abuse during childbirth, which is a violation of a universal human right that is due to every childbearing woman in every health system is common throughout the world and can occur at the level of interaction between the woman and provider, as well as through systemic failures at the health facility and health system level. Various studies across different countries have shown a high prevalence (physical abuse, non-consented care, non-confidential care, non-dignified care, discrimination based on specific client attributes, abandonment of care and detention in health care facilities)and negative impact of disrespect and abuse in facility-based childbirth on skilled care utilization [2, 12, 14–28].

In Ethiopia however, the few studies that have been conducted with regard to disrespect and abuse during childbirth in health facilities have revealed a high prevalence of the problem while we could not find any published research that is conducted in this study area[12, 16, 29]. This study was therefore, aimed at assessing the status of women’s disrespect and abuse during facility-based childbirth in Arba Minch town, south Ethiopia.

## Materials and methods

### Study area and period

This study was conducted in public health facilities in Arba Minch town. Arba Minch is the administrative town of Gamo Gofa zone which is one of the 14 zones in Southern Ethiopian Nations, Nationalities and Peoples Region with an estimated population of 2,043,668. Arba Minch town is located at a distance of 495 kilometers south from Addis Ababa. The total population of the town is about 79,961. There is one governmental hospital and two public health centers that are providing curative and preventive health services including maternity care to the community in and around the town. Arba Minch general hospital, the only hospital in the town has been serving as a referral hospital for patients from surrounding districts in the zone as well as from surrounding zones. The study period was from January 1 – February 28, 2017.

### Study design

Institution based cross-sectional study design was employed. Women who utilized one of the three public health facilities for childbirth purpose were included in the study. Women who underwent elective cesarean section were excluded from the study.

### Sample size and sampling technique

A single population proportion formula was used to calculate the minimum sample size required at 95% confidence level (CI), Z (1-ά/2) = 1.96), and level of non-respectful and abusive maternity care of 79%[29] and, 5% margin of error. This gives a sample size of 255 and with the addition of 10% for possible non response; the final sample size was 281.

There is one public hospital (Arba Minch general hospital) and two public health centers (Sikela and Shecha health centers) in Arbaminch town and all were included in the study. The average number of deliveries per month in *Arbaminch hospital, Sikela health center* and *Shecha health center* was 200, 60 and 30 respectively. During the two months of data collection period number of deliveries in the hospital was expected to be at least 400 while the health centers was estimated to be 120 at *Sikela* and 60 at *Shecha* which gives a total of 580 deliveries.

To include 281 in the study, proportional allocation method was used: 194 women from the hospital, 58 women from Sikela health center and 29 women from Shecha health center were interviewed by using systematic sampling technique by the assumption of: **N** (the average deliveries in two months period in the health institutions which is 580, and **n** (required minimum sample size = 281 which gives a k of 2): K = N/n =⇒ 580/281 (400/194, 120/58 or60/29) ≈ 2. To start data collection, the first two women from each health institution who had given birth at the health institutions on or after the first day of data collection were given numbers 1 or 2 and one of them from each health institution was selected by lottery method. Every other woman from each health institution was then included in the study starting from the women who were selected.

## Study variables

### Independent variables

*Socio-demographic characteristics*: Age, residence, income level, occupation, educational status

*Obstetric characteristics*: Parity, history of antenatal care follow-up, history of previous institutional delivery, place of birth, type of delivery

### Dependent variable

*Non respectful and abusive care:* Physical abuse, non-consented care, non-confidential care, non-dignified care discrimination based on specific clients’ attributes, abandonment of care and detention in health facilities

## Operational definitions

**Physical abuse: The presence of at least one of the following activities by the care provider on the client:** beating, threatening with beating, slapping, pinching, restraining or tying down during labor, cutting or suturing of episiotomy cuts or perineal tears without the use of anesthesia and the use of fundal pressure to fasten the delivery of the baby.

**Non-consented care:** The presence of at least one of the following: providers not giving women or her relatives proper information about medical procedures, not asking for women’s permission to conduct medical procedures such as cesarean sections, episiotomies, hysterectomies, blood transfusions, tubal ligation, augmentation of labor; and coercing into a medical procedures such as a cesarean section.

**Non-confidential care:** The presence of at least one of the following: giving birth in a public view without privacy barriers such as curtains; and having healthcare providers share sensitive clients’ information, such as HIV status, age, marital status, and medical history, in a way that other people who are not involved in their care can hear.

**Non-dignified care:** A report by the client about at least one of the following: intentional humiliation, blaming, rough treatment, scolding, shouting at, women not allowed to bring companion to the labor ward, and ordering to stop crying while they are in labor pain.

**Discrimination:** Discrimination based on specific client attributes like race, age, HIV/AIDS status, traditional beliefs and preferences, economic status, or educational

**Abandonment of care: If there is any of the following practices: leaving laboring woman alone, women** giving birth by themselves at health facilities, failure of care givers to monitor women in labor and intervene in life threatening conditions.

**Detention in facilities:** Detain the clients because of bills or damage to the property of the health care facility.

**Grand multipara:** A woman who has given birth 5 or more times.

**Night time Deliveries:** Deliveries that has occurred between 6:00 PM & 6:00 AM.

## Data collection and quality control

The data were collected by four midwife tutors. Questions on socio-demographic and obstetric characteristics of the study participants and details of the seven respectful maternity care rights were collected by face to face interview. The principal investigator had been supervising the data collectors closely during the entire data collection period. Two days training on how to carry out their duty was given for the data collectors ahead of starting data collection. Pretest was carried out on 5% of mothers prior to the actual study period to check the consistency of the questionnaire and the ability of the data collectors to carry out the duty. The questionnaire was then modified based on the pretest results. All the questionnaires were reviewed and checked for completeness everyday by the investigators. Epi info version 7.1.2.0 was used for data entry to reduce possible errors during data entry. Each and every questionnaire was crosschecked with the entered data and all observed errors were corrected.

### Data processing and analysis

All the questionnaires were checked for completeness, coded and entered in to Epi Info version 7.1.2.0 and then transported to SPSS version 24 software package for analysis. Descriptive statistics such as mean, percentage and standard deviation, were determined. Bi-variable logistic regression was done to determine the association between each independent variable and the outcome variables. Variables with p-value less than 0.25 in bi variable logistic regression were entered to multivariate logistic regression to adjust the effect of confounders on the outcome variables. The degree of association between dependent and independent variables was determined using the odds ratio with confidence interval of 95% and p-value of 0.05.

### Ethical considerations

Ethical clearance was obtained from Nech Sar Campus Ethical Review Committee of Arba Minch University. An official letter of cooperation was written by the College of Medicine and Health Sciences to Arba Minch General Hospital, Sikela and Shecha health centers. Verbal informed consent was obtained from each study participant and each study participant was informed about the objective of the study and confidentiality of the information she was giving. The participants were also informed that they have full right not to participate in the study or to stop participation at any time during the interview.

## Results

### Socio-demographic characteristics of the study participants

A total of 281 women who had given birth at the governmental health institutions in Arba Minch town were interviewed making a response rate of 100%. The mean age of the study participants was 28.5 years(SD + 5.92). The majority (57.3%) of the women were house wives, from Gamo ethnic group (48.4%), Orthodox religion flowerers (49.8%), urban residents (64.8%), in marital relationship (98.6%) and did not have their own income (65.1%) **((Table1)**.

**Table 1.** Socio-demographic characteristics of the study participants, Arba Minch town, south Ethiopia, May, 2017 (n=281).

| Variable | Frequency | Percent |
| --- | --- | --- |
| **Age** |  |  |
| < 20 years | 12 | 4.3 |
| 21 – 34 years | 214 | 76.2 |
| ≥ 35 years | 55 | 19.6 |
| **Religion** |  |  |
| Orthodox | 140 | 49.8 |
| Protestant | 112 | 39.9 |
| Muslim | 13 | 4.6 |
| Others* | 16 | 5.7 |
| **Residence** |  |  |
| Rural | 99 | 35.2 |
| Urban | 182 | 64.8 |
| **Ethnicity** |  |  |
| Amhara | 19 | 6.8 |
| Gamo | 136 | 48.4 |
| Gofa | 41 | 14.6 |
| Konso/Derashe/Alle | 13 | 4.6 |
| Oromo | 17 | 6.0 |
| Wolita | 44 | 15.7 |
| Others** | 11 | 3.9 |
| **Marital Status** |  |  |
| Married | 277 | 98.6 |
| Divorced | 1 | 0.4 |
| Widowed | 3 | 1.1 |
| **Occupation** |  |  |
| Daily laborer | 15 | 5.3 |
| House wife | 161 | 57.3 |
| Student | 22 | 7.8 |
| Private Business | 40 | 12.2 |
| Governmental employee | 43 | 15.3 |
| **Educational status** |  |  |
| Unable to read write | 56 | 19.9 |
| Read & write | 28 | 10.0 |
| Elementary school (Grade 1 – 4) | 51 | 18.1 |
| Secondary school(Grade 5 – 8) | 72 | 25.6 |
| High /Preparatory school(Grade 9 – 12) | 25 | 8.9 |
| Above grade 12 | 49 | 17.4 |

\* Traditional, Catholic and & Jehovah Witnesses \*\* Guragie, Tigre & Amaro

### Obstetric characteristics of the study participants

More than 95% of the study participants had history of ANC follow up during the recent pregnancy. Almost two-third of them had history of previous institutional delivery and 214(76.2%) had given birth via vaginal route. Two hundred eight (74.0%) of the women’s preferred birthing position was kneeling **(Table 2)**.

**Table 2.** Obstetrics characteristics of the study participants, Arba Minch town, south Ethiopia, May, 2017 (n=281).

| Variable | Frequency | Percent |
| --- | --- | --- |
| **Place of the index delivery (n= 281)** |  |  |
| Arba Minch general hospital | 194 | 69.0 |
| Shecha HC | 29 | 10.3 |
| Sikela HC | 58 | 20.7 |
| **History of ANC Follow up (n = 281)** |  |  |
| Yes | 268 | 95.4 |
| No | 13 | 4.6 |
| **Parity (n= 281)** |  |  |
| Primiparous | 74 | 26.3 |
| Multiparous | 167 | 59.4 |
| Grand multiparous | 40 | 14.2 |
| **History of previous institutional delivery (n= 281)** |  |  |
| Yes | 177 | 63.0 |
| No | 104 | 37.0 |
| **Route/Type of current delivery (n = 281)** |  |  |
| Vaginal | 214 | 76.2 |
| Cesarean | 67 | 23.8 |
| **Time of delivery (n= 281)** |  |  |
| Day | 135 | 48.0 |
| Night | 146 | 52.0 |
| **Number of birth attendants (n= 214)** |  |  |
| 1 - 2 | 121 | 56.5 |
| 3 - 4 | 74 | 34.6 |
| 5 - 8 | 19 | 8.9 |
| **Sex of the main birth attendant (n= 214)** |  |  |
| Female | 153 | 71.5 |
| Male | 61 | 28.5 |
| **Preferred birthing position (n= 281)** |  |  |
| Kneeling | 208 | 74.0 |
| Laying on back /Lithotomy | 67 | 23.8 |
| Squatting | 6 | 2.1 |
| **Labor was augmented/induced (n = 281)** |  |  |
| Yes | 30 | 89.3 |
| No | 251 | 10.7 |
| **Transfused with blood** |  |  |
| Yes | 1 | 0.4 |
| No | 280 | 99.6 |

### The overall prevalence of non-respectful and abusive care

While the seven women’s rights were not equally violated; ranging from zero percent in the seventh right: the women’s right to liberty, autonomy, self-determination, and freedom from coercion to more than 90% in the second right: the women’s right to information, informed consent, overall 278(98.9%) of the women have reported that they had faced at least one of the violation of their seven rights. The next sections describe violations of the individual rights.

### Violation of article 1: the women’s right to be free from harm and ill treatment (prevalence of physical abuse)

Two women (0.7) % have reported that the birth attendants have used physical forces (slapping) against them while they were in labor pain while 4(1.4%) of the women said that the birth attendant have threatened them with use of physical force. Among those women who gave birth vaginally, 64 had their premium sutured as a result of tear or after episiotomy. Among them 12(18.8%) have reported that their perineum was sutured without the use of any anesthesia, 35(16.4%) have said that the birth attendants have pushed their abdomen down (used fundal pressure) to deliver their babies. One hundred thirty six (63.6%) of the women who gave birth vaginally declared that they were not allowed to adopt the position of their choice to bear down. Regarding ambulation, 87(31%) of the women were not allowed to ambulate before giving birth of which 26(29.9%) were not told reasons for restricting ambulation while 21(7.5%) of the women (18 of these (85.7%) finally underwent cesarean reported that they were restricted from any fluid during the course of labor. There was no woman who reported that her legs were tied down by strips during labour or her family was forced to clean the delivery bed or delivery room. Overall, the prevalence of physical abuse was 29.5% (95% CI: 24.2, 34.8).

### Factors associated with physical abuse

In bi-variable analysis: being rural resident, having no ANC follow up during the index pregnancy, having no previous history of institutional delivery, and giving birth at hospital were factors associated with physical abuse. However, multivariate analysis demonstrated that being rural resident and delivery in the hospital were the only factors which were significantly associated with physical abuse.

Women who gave birth at hospital were about 2 times more likely to be physically abused compared with those who gave birth at health centers (AOR = 2.82, 95% CI: 1.48- 5.35). Those women who were from rural areas were about 2 times more likely to face physical abuse than their urban counterparts (AOR = 1.85, 95% CI: 1.07 – 3.18) (**Table3**).

**Table 3.** Bi variable and multivariate analysis of factors associated with physical abuse during childbirth at health institutions in Arba Minch town, south Ethiopia, May, 2017 (n=281).

| Variables | Physical abuse |  | OR (95% CI) |  |
| --- | --- | --- | --- | --- |
|  | Yes | No | COR (95% CI) | AOR (95% CI) |
| **Type of health institution** |  |  |  |  |
| Hospital | 68(35.1 %) | 126 (64.9%) | 2.59(1.38,4.86) | **2.82 (1.48, 5.35)** |
| HC | 15(17.2 %) | 72(82.8%) | 1 | **1** |
| **Residence** |  |  |  |  |
| Rural | 36(36.4%) | 63(63.6%) | 1.64(0.96, 2.78) | **1.84(1.07, 3.18)** |
| Urban | 47(25.8%) | 135(74.2%) | 1 | **1** |
| **ANC follow up** |  |  |  |  |
| No | 6(46.1%) | 7 (53.9%) | 2.13 (0.69, 6.53) | * |
| Yes | 77(28.7%) | 191(71.3%) | 1 |  |
| **History of previous institutional delivery** |  |  |  |  |
| No | 38(36.5 %) | 66(64.5%) | 1.69 (1.00, 2.85) | * |
| Yes | 45(25.4%) | 132(74.6%) | 1 |  |

Hospital: P value: 0.02, Rural residence: P value = 0.027 \*= Not significant in multivariate analysis

### Violation of article 2: women’s right to information, informed consent (prevalence of non-consented care)

The women’s right to information and informed consent was the most frequently violated right as reported by the study participants. Only 19(6.8%) of the study participants have said that the care givers introduce themselves during the admission time while about 208(74%) reported that they were not told about the evaluation of their initial assessment by the care givers. During the initial assessment, only 57(20.3%) of the respondents said that they were encouraged to ask about unclear points. Among the 53 women for whom episiotomy was performed, only 21(39.6%) had provided verbal consent. Out of the 67 women who underwent cesarean birth 61(91%) reported that they were well informed before the procedure while 17 (56.7%) of the 30 women whose labour was augmented were told about the procedure in advance. Overall, 260(92.5%, 95%CI: 90.9, 94.1) of the respondents reported that their consent was not asked at one of the events.

### Violation of article 3: women’s right to privacy and confidentiality (prevalence of non-confidential care)

Women’s right to privacy and confidentiality is among the three women’s right which were less violated. Two hundred fifty eight(91.8%) of the women responded that, no one except those health care providers who were involved in their birth attendance entered the delivery room while they were naked. Almost all (97.2%) of the respondents have not heard the care providers sharing their secret information to others or they trust the care providers that they would not share their secret. The majority, 255 (90.7%) of the respondents were satisfied by the physical barriers or the use of curtains to keep their privacy during the course of labour and delivery. Overall, 233(82.9%) of the respondents reported that their care was confidential which makes the prevalence of non-confidential care of 17.1% (95% CI: 12.7, 21.5).

### Factors associated with non-confidential care

Being rural resident and giving birth at hospital were factors which are significantly association with non-confidential care. Women who gave birth at hospital were about 3 times more likely to report non-confidential care when compared to those who gave birth at health centers (AOR = 3.34, 95% CI: 1.42, 7.87). Those women who were from rural areas were about 2 times more likely to report non-confidential care when compared to their urban counterparts (AOR = 1.93, 95% CI: 1.013, 3.68)(**Table 4**)

**Table 4.**
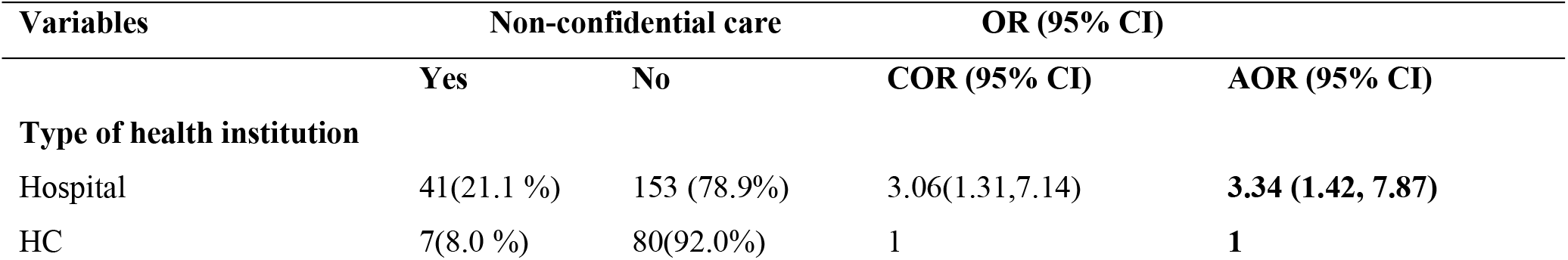
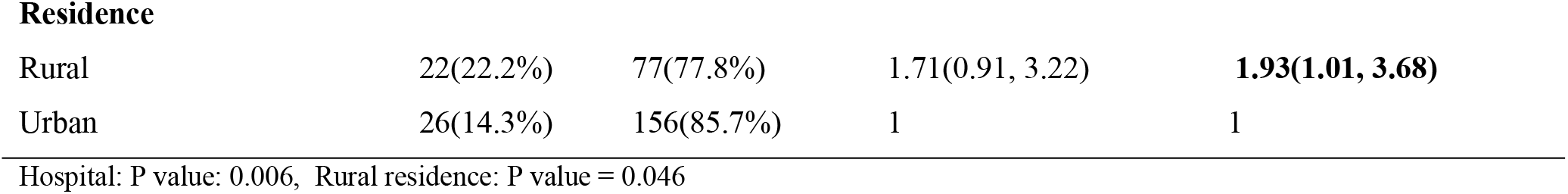
Bi variable and multivariate analysis of factors associated with non-confidential during childbirth at health institutions in Arba Minch town, south Ethiopia, May, 2017 (n=281).

| Variables | Non-confidential care |  | OR (95% CI) |  |
| --- | --- | --- | --- | --- |
|  | Yes | No | COR (95% CI) | AOR (95% CI) |
| **Type of health institution** |  |  |  |  |
| Hospital | 41(21.1 %) | 153 (78.9%) | 3.06(1.31,7.14) | **3.34 (1.42, 7.87)** |
| HC | 7(8.0 %) | 80(92.0%) | 1 | **1** |
| Residence |  |  |  |  |
| Rural | 22(22.2%) | 77(77.8%) | 1.71(0.91, 3.22) | **1.93(1.01, 3.68)** |
| Urban | 26(14.3%) | 156(85.7%) | 1 | 1 |

Hospital: P value: 0.006, Rural residence: P value = 0.046

### Violation of article 4: women’s right to be treated with dignity and respect (prevalence of non-dignified care)

Nighty seven (34.5%) of the respondents reported that the health care providers were not talking to them politely while 8(2.8%) of them said that the care providers blame them for getting pregnant. About a quarter (23.1%) of the participants said that the birth attendants have shouted at them to calm them down while they were in severe labour pain. Nearly half of the respondents responded that, their relatives were not allowed to accompany them during the course of labour. Overall, 104 women reported at least one form of the violation of this right making the overall the prevalence of non-dignified care to be 36.7% (95% CI: 34.9–38.5).

### Factors associated with non-dignified care

In bi-variable analysis, low educational status or having no formal education, recent delivery by cesarean section and giving birth at hospital were factors which were associated with non-dignified care. However, multivariate analysis has showed that births attended at hospital and having no formal education or less grade schooling were the only factors which were significantly associated with non-dignified care.

Women who gave birth at hospital were about 10 times more likely to face non-dignified care compared with those who gave birth at health centers (AOR = 9.93, 95% CI: 4.37, 19.76). Those women who have no formal education were about 3 times more likely to be treated in a non-dignified way when compared to those who have at least completed secondary school (AOR = 3.17, 95% CI: 1.55, 6.49). (**Table 5**).

**Table 5.**
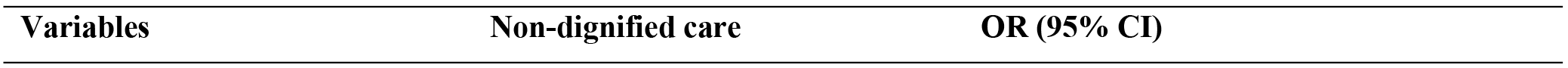
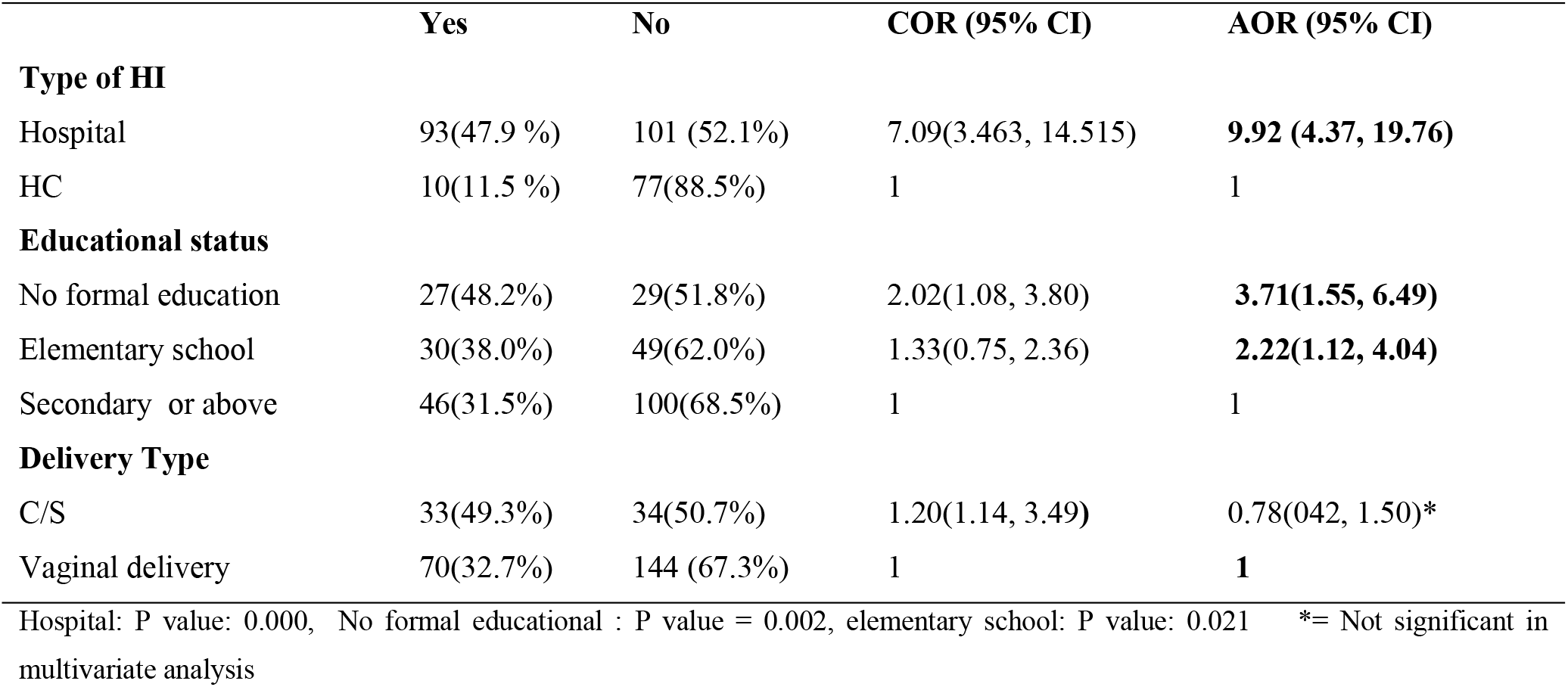
Bi-variable and multivariate analysis of factors associated with non-dignified care during childbirth at health institutions in Arba Minch town, south Ethiopia, May, 2017 (n=281).

|  | Yes | No | COR (95% CI) | AOR (95% CI) |
| --- | --- | --- | --- | --- |
| **Type of HI** |  |  |  |  |
| Hospital | 93(47.9 %) | 101 (52.1%) | 7.09(3.463, 14.515) | **9.92 (4.37, 19.76)** |
| HC | 10(11.5 %) | 77(88.5%) | 1 | 1 |
| **Educational status** |  |  |  |  |
| No formal education | 27(48.2%) | 29(51.8%) | 2.02(1.08, 3.80) | **3.71(1.55, 6.49)** |
| Elementary school | 30(38.0%) | 49(62.0%) | 1.33(0.75, 2.36) | **2.22(1.12, 4.04)** |
| Secondary or above | 46(31.5%) | 100(68.5%) | 1 | 1 |
| **Delivery Type** |  |  |  |  |
| C/S | 33(49.3%) | 34(50.7%) | 1.20(1.14, 3.49) | 0.78(0.42, 1.50)* |
| Vaginal delivery | 70(32.7%) | 144 (67.3%) | 1 | **1** |

Hospital: P value: 0.000, No formal educational : P value = 0.002, elementary school: P value: 0.021 \*= Not significant in multivariate analysis

### Violation of article 5: women’s right to equality, freedom from discrimination, and equitable care (prevalence of discrimination based on specific client attributes)

While no woman was discriminated because of her religion or because her retroviral infection status, 2 (0.7%) women perceived that the birth attendants have discriminated them because of their traditional beliefs. Thirty five (12.5%) of the respondents said that they were discriminated (perceived) because they were from rural area or because their educational status is low or have no formal education. Another 4 (1.4%) of the women have reported that they were discriminated because they were too young to give birth. Overall, 51 of the respondents felt discriminated because of their different attributes making the prevalence of discrimination 18.1% (95% CI: 13.6, 22.6).

### Factors associated with discrimination

Education status of the respondents, residence, place of delivery, type of delivery, history of previous health institutional delivery and occupation of the respondents were factors which were associated with discriminating care in bi-variable logistic regression analysis. In multivariate logistic regression analysis; place of delivery, residence, history of previous health institution delivery and educational status of the respondents were factors which were significantly associated with discriminated care during childbirth.

Discrimination based on some women’s attributes was about 9 times more common in hospital than in health centers (AOR = 9.14, 95% CI: 3.17, 26.34). Those women who have no formal education were about 15 times more likely to felt discriminated when compared to those who have at least completed secondary school (AOR = 14.82, 95% CI: 5.22, 39.25). Women who were from rural areas were about 4 times more likely to felt discriminated than those who are from urban area (AOR = 3.76, 95% CI: 1.70, 8.31). Those women who have no previous history of health institutional delivery were about 4 time more likely to felt (perceive) that they were discriminated than those who have previous history of institutional birth (AOR = 4.44, 95% CI: 1.97, 9.10) (**Table 6**).

**Table 6.**
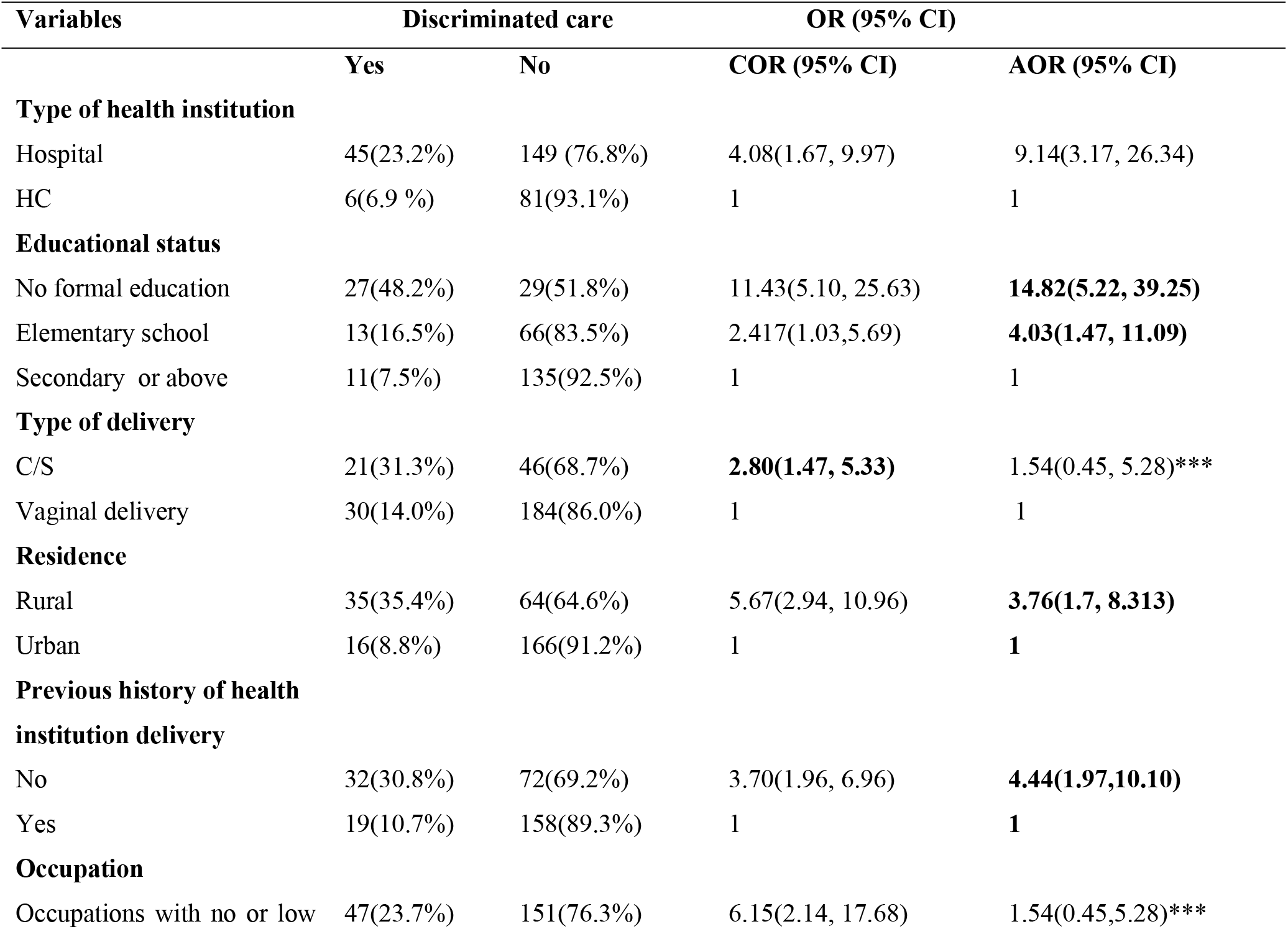
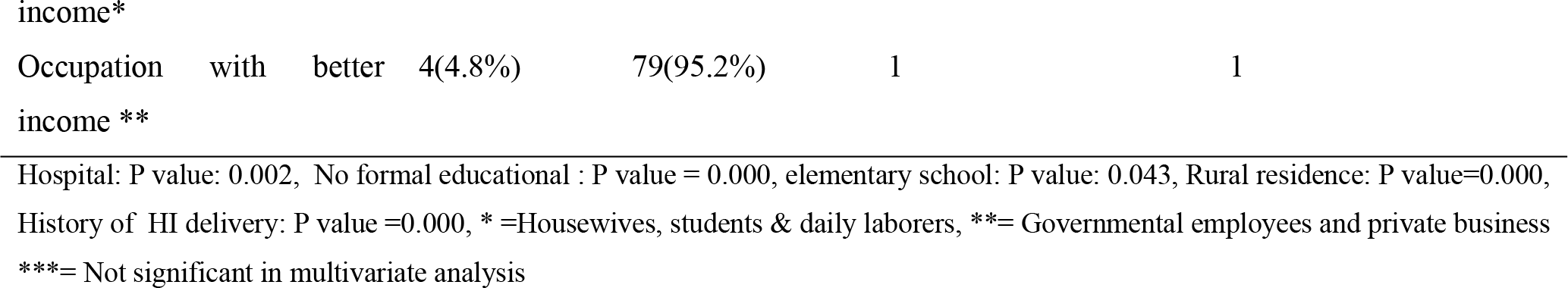
Bi-variable and multivariate analysis of factors associated with discriminated (actual/perceived) care during childbirth at health institutions in Arba Minch town, south Ethiopia, May, 2017 (n=281).

### Violation of article 6: women’s right to healthcare and to the highest attainable level of health and continuous support (prevalence of abandonment of care)

Abandonment of care was the least to be reported by the study participants next to detention in health facilities. Only 12(4.3%) of the study participants responded that they were left alone for some period of time while they were in need of someone to be with them. However, none of the participants said that they had given birth by themselves in the health facilities because the health care providers were not around or no health care provider reached them when they have encountered life threatening condition and shouted for help. The overall prevalence of abandonment of care is therefore 4.3%, 95% CI: 3.1, 5.5).

### Violation of article 7: Women’s right to liberty, autonomy, self-determination, and freedom from coercion (prevalence of detention in health facilities)

From this study no study participant responded that she or her families were detained in the health facilities for the issue of payment or damage to the health institutions’ equipment.

## Discussion

This study has investigated the status of disrespect and abuse during childbirth using the seven universal rights of child bearing women[1, 2]. Almost all (98.9%) of the women have reported that they have faced at least one form of disrespect or abuse during their stay at the health care facilities.

This figure is similar with findings from one Nigerian study which was carried out in 2012 where 98% of the women had reported at least one form of disrespect[30]. However, it is higher than findings from a study conducted in Addis Ababa in Ethiopia in 2013 where the overall prevalence of disrespect and abuse was 76.8%[29]. The higher prevalence in this study in comparison to the previous Ethiopian study could be because of the differences in verification criteria as we have included the use of obsolete procedures which put women’s health at risk like the use of fundal pressure to expell babies and suturing episiotomies or perineal tears without use of local anesthesia as physical abuse. It could also be explained by the differences in study settings as the previous study was conducted in the capital Addis Ababa, the capital city of the country, where the status of clients is expected to be higher. This too high prevalence of abuse and disrespect in studies suggests “normalization of disrespectful care” by clients as well as by health care providers in low income countries[2, 28].

As the prevalence of violation of the seven women’s right ranged from zero (detention in health facilities) to nearly 100% (non-consented care), we will discuss the prevalence and factors associated with each disrespects separately as it will have different implications.

In this study the prevalence of physical harm and ill-treatment was 29.5% (95% CI: 24.2, 34.8) which is preceded only by non-consented care. This is in line with the findings from a study conducted in Nigeria in 2012 where the prevalence of physical abuse was 35.7% and the previous Ethiopian study where it was 32.9%[29, 30]. From this we can learn that, physical abuse has been persistently high since the last 5 years. However, as groups and criteria are not comparable no statements on evolution in the prevalence can be made.

Two mothers (0.7%) (both from the hospital) have reported that the birth attendants have beaten them while they were in labour pain. This is in line with the results from a previous study conducted in 5 East and South African countries from 2009 to 2012 where the prevalence of the use of physical force like slapping was 0.83%[12]. However, it is lower than the findings from a study conducted in Addis Ababa Ethiopia in 2013 where physical force was used in 2.3% of the mothers[29].

The use of fundal pressure to deliver babies during second stage of labour was reported by 16.4% of the 214 women who had given birth via vaginal route, in this study. This is also concurrent with the study conducted in 5 countries including Ethiopia in East & Southern Africa where Ethiopia was the only country where fundal pressure to aid delivery of babies was reported[12]. This makes Ethiopia, the country where some of the obsolete procedures are still practiced by some health care providers.

Not using local anesthesia for episiotomy or perineal tear repair was reported by 12(18.8%) women out of the 64 women whose perineum was sutured. This is also reported by the study conducted in 5 countries where anesthesia for episiotomy or perineal tear repair was not used by care providers in Ethiopia[12]. Even though there is an improvement in the use of local anesthesia from not using at all to more than 80%, it should be practiced in all of the laboring mothers as it is inhuman, unprofessional and also contributes to non-preference of health institution deliveries[2].

Forty eight (81.3%) of the 59 primiparous who had given birth vaginally had episiotomy. Even though the use of episiotomy by itself is not a physical abuse, routine episiotomies can result in unnecessary perineal scars which might lead to dyspareunia.

Overall, women who gave birth at hospital were about 3 times more likely to be physically abused compared with those who gave birth at health centers. This is consistent with the previous study conducted in Addis Ababa and [29]. This could be attributed to workload and dissatisfaction by the hospital staffs, presence of a different mix of health professionals with different backgrounds, client’s inability to communicate with the staffs due to language barriers (for those who are referred from rural health centers).

Moreover, those women who were from rural areas were about 2 times more likely to face physical abuse than their urban counterparts. This is against their right as humans and will result in increased homebirths which are attended by unskilled persons in the future[2]. Obviously, this will pose a great challenge for the improvement of the maternal health of the country as more than 85% of Ethiopian population resides in rural areas.

However, no mother has reported that her legs were tied down during childbirth which indicates that some of the obsolete procedures have been abandoned as it used to be practiced routinely [2].

Non-consented care or the women’s right to information, informed consent, and preferences was the most frequently violated women right in Ethiopia as more than 92 % of the women have reported it in this as well as in previous study conducted in Ethiopia[29]. A study from Nigeria revealed that the prevalence of non–consented care was 45.5% which is less than half of the prevalence in Ethiopia[30]. This indicates that the right to information and consent is not regarded as a right by Ethiopian health care providers when compared to other women’s rights.

While only 9% of the 67 women who underwent cesarean section have said that they were not well informed about the procedure beforehand, the proportion of women who were consented before induction or augmentation of their labour was 43.3% and the figure is higher in case of performing episiotomy which is 60.4%. Not taking consent before procedures was also reported in 38% in the multi country study and 48% in previous study in Ethiopia[12, 29]. From this high figure, we can learn that the health care providers are taking consent not because it is the right of the client but to be on the safe side in case if something goes wrong as it is very high before cesarean section; the procedure that carries major risk and very low before episiotomy; a procedure with relatively lower risk.

The above interpretation can be supported by the result that very high proportion (93.2% in this study and 89.0%in the previous Ethiopian study[29] of women reported that the health care providers have not introduced themselves at admission. Telling the findings of their initial assessment which will have high potential to prepare the clients on what to expect was not done in 26% of the respondents which is relatively lower than the multi country study finding which was 33%[12]. What is being done and what to expect was not told to 57% of the women in this study and 48% of the early study[29]. If women do not take an active role in their healthcare decisions, they will not develop a better understanding of their choices and will less likely receive care consistent with their preferences, values, and goals. This might negatively affect skilled birth attendance in the future [2].

Regarding non-confidential care, a small proportion of women (17.1%) responded that their privacy was not kept during the course of labour and delivery. This is relatively lower than the finding from previous multi country study [Zanzibar (78%), Ethiopia (73%), Tanzania (46%), Kenya (35%), Madagascar (28%) and Rwanda (22%)[12]. But it is comparable with the results from another study conducted in Ethiopia where the prevalence was 33% in hospitals and 9% in health centers[29]. It is also lower than the prevalence of non-consented care and physical abuse mentioned above in this study.

This could be because it is a factor which can be modified easily. For instance physical barriers like curtains can be arranged by the administrators of the health institutions unlike that of human behavior which may take a longer period to change. It could also be because the number of health institutions participated in this study is very small and the presences of physical barriers like curtains mean non-or all for the health institution although the clients may report the inappropriate use of curtains during her course of labour.

Women who gave birth at hospital were about 3 times more likely to report non-confidential care when compared to those who gave birth at health centers. This supports the evidences from previous studies[29]. The high prevalence of non-confidential care at hospital can be explained by the fact that hospital staffs are carrying for large number of clients at a time, the occurrence of emergency condition which sometimes necessities the introduction of new persons to the rooms and exposure of nearby clients for the emergency treatment of the neighbor client. Those women who were from rural areas were about 2 times more likely to report non-confidential care when compared to their urban counterparts. This can be related to the violation of the second article as the clients are not informed about anything about the course of their labour and will have significant effect on the next institutional deliveries[2].

The prevalence of non-dignified care in this study is relatively higher than the findings of the similar previous studies. While it is 36.7% in this study, it was between 10% to 20% in the previous studies[12,29]. Shouting at the clients while they were screaming because of labour pain was reported by 23.1% of the respondents while it was less than 9% in one study conducted in Tanzania[11]. However, the prevalence of blaming the women for getting pregnant and the use of rude words and insulting was reported by only 2.8% of the respondents which is lower than the results from the previous Ethiopian study where it was 7.5%[29]

The reason for the higher prevalence of non-dignified care in this study could be the inclusion of not allowing companion to the delivery room in to this category while it is usually categorized under women’s preference section or abandonment of care. As it is known those articles usually overlap and are non-exclusive and here we took this under the non-dignified care because women feel more hopeless, when they become alone.

Those women who have no formal education were about 3 times more likely to be treated in a non-dignified way when compared to those who have at least completed secondary school. This could be related to the violation of the right of the women to information. For instance, if a woman is told about the reasons why she should be alone in case of procedures which may not be comfortable for the relatives, the client might not feel this separation from her families as an issue of dignity.

Women who gave birth at hospital were about 10 times more likely to face non-dignified care compared with those who gave birth at health centers. This can be explained by the same reasons mentioned under the previous sections.

In this study about 18% of the responds perceived as if the care providers discriminated them because of their status which is also reported in other countries like Peru, Burkina Faso and Kenya as it was stated in hills landscape analysis[2].

Discrimination based on some women’s attributes was about 9 times more common in hospital than in health centers. Those women who have no formal education were about 15 times more likely to perceive as if they were discriminated when compared to those who have at least completed secondary school. Women who were from rural areas were about 4 times more likely to report discrimination than those who are from urban area. Those women who have no previous history of health institutional delivery were about 4 times more likely to feel that they were discriminated than those who have previous history of institutional birth.

All the factors associated the perceived or real discrimination could be because in the hospital many women are admitted at a time and those who are new for the environment or have difficulty of coping with the environment can easily perceive as if they were discriminated. Sometimes not knowing the nature of labour may lead to perceiving being discriminated. For instance, if a woman from rural area with early stage labour comes first and an educated urban woman comes with an advanced labour and the staff manages the urban women first, the uneducated rural women may consider this as discrimination. The main point is, be it real or perceived, the consequences will be negative on the future utilization of health institution delivery services and hence need to be treated equally. This can also be linked to the client’s right to information.

Even though the sixth article of rights of child bearing women sates that every woman has the right to healthcare and to the highest attainable level of health[1], women in labour pain are sometimes left alone. About 17% of the participants of this study responded that they had been left alone at least once for some period of time even though there was no mother who said that she had given birth by herself in the health institutions. This result is relatively promising when compared to the previous studies where about 63.6% of the women who had given birth at hospitals in Addis Abba responded that they were abounded during childbirth. But the figure is still high when compared to other similar study in other African countries like Tanzania where it was 7.9%[11, 29].

## Conclusions and recommendations

The status of non-respectful care in the health care facilities in this study area was very high especially in the hospital. The problem is particularly high among uneducated women and women who are from rural areas. Women’s right to information and informed consent was the most frequently violated while detention in health care facilities was reported by none of the respondents. Still there are practice of some obsolete procedures like use of fundal pressure to aid explosion of the baby, unnecessary episiotomies, and nonuse of anesthesia to suture perineum which cane physically harm the women.

This unacceptably high prevalence of non-respectful maternity care at health care facilities needs critical attention by the health managers. Especially the health care providers should be oriented on the importance of the informed consent and to avoid obsolete procedure appropriate management of episiotomies and perineal tears. Some of the misconducts might be related dissatisfaction of staffs and also needs serious attention by the health managers of the institutions.

## Acknowledgments

We would like to acknowledge the study participants for their cooperation by sacrificing their time and proving information necessary for this study. We are also grateful to Professor Dr. Yves Jacquemyn for reviewing the final manuscript.

## Author contributions

Conceived and designed the research work: GGU MKG WGB. Analyzed the data: GGU MKG WGB Wrote the paper: GGU MKG WGB. Read and approved the final manuscript; GGU MKG WGB.

